# Adapting Indirect Calorimetry to measure metabolic status of healthy and septic neonatal mice

**DOI:** 10.1101/2024.05.23.595520

**Authors:** Adrien Eynaud, Joan Solomon, Elizna Schoeman, Ziyuan Wu, Nelly Amenyogbe

## Abstract

Commercially available platforms to measure murine pulmonary gas exchange have long been used to measure metabolic status of adult animals, thus providing insights into metabolic disease, diabetes, and infection. Metabolic status is increasingly being recognized as an important modulator of neonatal immunity, and capturing pulmonary gas exchange in neonatal animals provides a non-invasive way to capture physiological information in health and disease and may reveal metabolic determinants of immune-mediated diseases unique to this life stage. We evaluated an indirect calorimetry (IC) system, the Promethion Core CGF system outfitted with Respirometry Chambers (RC3) as a tool to accurately capture pulmonary gas exchange from individual healthy and septic murine neonatal pups in the first week of life. We assessed the technical variance of the platform, impact of the procedure of animal welfare, compared measurements performed either at room temperature or at 30°C, and determined the platform’s sensitivity to measure gas exchange from pups with very small lung capacity or low respiratory rate. While gas exchange was not captured above background levels in all pups with either very small lung volume (pups less than 4 days old) or with very low respiratory rates (septic pups with the most depressed respiratory status), measurements did capture physiologically relevant changes in gas exchange across age and disease states. The impost associated with frequent handling of septic animals for IC did not negatively impact clinical outcomes among pups challenged with a polymicrobial slurry. Further, while performing the IC readings at 30°C successfully stabilized animal body temperature, the VO2 and VCO2 values differed across temperature states for older pups. In conclusion, the Promethion Core system outfitted with RC3 chambers is a viable platform to integrate IC into murine neonatal health research.

## Introduction

The role of host metabolism in determining the quality of immune responses to infection is increasingly being recognized (*1, 2*). While experimental reductionist measures of immunometabolism most often focus on metabolic pathways at the single-cell level, the importance of holistic measures of whole-body metabolism as a central element in orchestrating the immune response is emerging as critical (*2, 3*). Preclinical models of infection have largely relied on bio-sampling post-mortem to measure biologically relevant indicators of physiological response, including immunity and metabolic status. This is especially true for neonatal mice, where there are no humane and robust methods to collect relevant biological specimens (e.g. blood; liver) longitudinally, as even minor surgeries result in most neonatal mice being cannibalized by the dam (*4*). This results in a substantial disadvantage, as measurements taken at early courses of infectious processes can be the most informative; but if animals are euthanized at those time points, then connecting early or baseline physiological responses to long-term disease severity or survival becomes impossible. Indirect calorimetry provides a solution.

Indirect calorimetry (IC) allows for non-invasive, continuous measurement of host metabolic status by capturing the amount of Oxygen (O_2_) consumed and amount of Carbon Dioxide (CO_2_) released by an organism – pulmonary gas exchange. With these two measures, IC is able to convert rates of gas exchange to calculate a diverse set of physiologically relevant measures; velocity of O_2_ (VO_2_) and CO_2_ (VCO_2_) gas exchange, energetic expulsion in kilocalories per hr per gram body weight (REE), and the respiratory exchange ratio (RER) (*5*). IC has been employed to non-invasively study animal models of health and disease including healthy ageing (*6, 7*), obesity (*8, 9*), diabetes (*10*), links between the gut microbiome and metabolism (*11*), and exercise biology (*12*). IC has also recently been incorporated into studies of the host response to infection in adult animals, with the landmark finding that the host re-directs energy from maintaining homeothermy to resolve inflammation. This is also reflected by a decrease in REE during inflammation driven by extracellular, but not intracellular infection (*13*). IC investigations in neonatal mice are rare, and have been used only to measure short-term oxygen consumption across a range of atmospheric temperatures (*14*).

The Promethion Core system (Sable Systems) for mice is designed to apply indirect calorimetry (IC) *in vivo* to living preclinical models by measuring respiratory gas exchange (VO_2_ and VCO_2_), and from those values, determining the respiratory exchange ratio and energetic expenditure (EE) of experimental animals. Respirometry chambers (RC) conveniently capture pulmonary gas exchange from adult animals via cyclic continuous measurements. However, this system is impractical to capture metabolic status of individual neonatal pups, who cannot be individually housed, and with much smaller lung volumes than adult rodents for which these cages were optimized. Individual measurements are indispensable, as variance in clinical outcome may be associated with distinct metabolic adaptations to infection that are lost when measurements are performed on the level of an entire litter. We therefore developed a protocol to apply 700 mL glass cylindrical RC3 chambers to longitudinally measure the pulmonary gas exchange among the same neonatal pups at intermittent time points between day of life (DOL) 1 and 7. We determined whether cross-sectional measurements taken from the same pups across two cycles provided reliable data points, and whether the chambers offered enough sensitivity to capture gas exchange among pups with either very low pulmonary volume (pups younger than 4 days of life) or septic pups in a hypometabolic and depressed respiratory state. We also determined whether the stress and extra handling associated with collecting IC measurements frequently from septic pups negatively impacted their disease course, as this would then represent a significant confounder to assessing the impact of interventions Lastly, we determined the impact of conducting readings at 30°C to minimise body-heat related VO2 and VCO2 variations, comparing these results to measurements taken at room temperature in healthy and septic pups. Overall, we found that the Promethion Core outfitted with RC3 chambers is a humane method to reliably capture and longitudinally follow the metabolic status of individual neonatal pups.

## Methods

### Animals

Specific pathogen-free C57BL/6J mouse pups were bred in Telethon Kids Institute housing facility and housed under standard conditions of 12 h light and 12 h dark cycles. Pregnant females were separated from males and then monitored daily (day-time hours) thereafter to ensure accurate identification of date of birth. Litters varied in size from four to eight pups. All animal procedures were approved by the Telethon Kids Institute’s Animal Ethics Committee under protocol numbers AEC 370 and AEC 363. All procedures performed were in accordance with institutional and national guidelines.

### Polymicrobial sepsis

We have previously standardized and described a neonatal mouse model of neonatal polymicrobial sepsis (*15-17*). Briefly, a homogeneous cecal slurry stock is prepared by pooling cecal contents from 7-10 week-old C57BL/7 mice, diluting the slurry in 5% dextrose water, and freezing in aliquots at -80°C. Polymicrobial sepsis is induced by injecting a weight-adjusted dose of cecal slurry via intraperitoneal injection on DOL 7-8. Pups are observed for any injection-related complications at 2 hours post challenge (hpc) and then monitored for sepsis-related morbidity and the identification of mice at a humane endpoint. Pups are monitored for clinical signs by removing the dam from the nest, and measuring the animal’s body weight and behavioural score as previously described (*15, 16*). Body temperature was measured using the Bioseb Rodent Thermometer (BIP-IRB153) by scruffing the neonatal mouse, and steadily holding the probe over the trunk of the mouse for 5-10 seconds. This method is only suitable before neonatal mice grow fur on the ventral side, at approximately DOL 10.

### Indirect calorimetry measurements via the Promethion Core system

The Promethion Core™ CGF system (Sable, Croydon) was used to perform IC measurements. Our custom system setup comprised eight 700 mL RC-3 inline respirometry glass chambers (Sable, Croydon) containing a stainless-steel internal platform. The system is equipped with a baseline container to measure the oxygen and carbon dioxide content in the room. To measure pulmonary gas exchange from neonatal animals, the dam was first removed from the nest, and pups were monitored for weight and clinical signs using the same procedures applied to monitor mice following septic challenge. After monitoring was performed for all pups, one pup was placed within each of eight RC3 chambers. A cardboard tube, lined with a soft wipe, was placed on the internal platform to shield pups from light and to avoid placing pups on a cold metal surface (Figure 1A). When capturing experimental data, the Promethion Core first captures data from the baseline chamber and then cycles through each of the eight RC3 chambers using 30 second cycles with a flow rate of 2000 mL/min. With this, the Promethion Core requires approximately 3 minutes to complete one reading from each chamber. Given that our use of RC3 chambers requires pups to be completely isolated from the dam and their littermates, our protocol involves cycling through each RC3 chamber twice to capture two readings during each cross-sectional time point. These readings were treated as replicates and averaged to generate one value per animal per time point. Pups were returned to the nest with dam immediately after completion of measurements for all animals. For each experiment, pups within each litter were assigned to one RC3 chamber. If the pup was euthanized, having reached humane endpoint, then that chamber remained empty for the duration of the experiment. Using this method, we accumulated 29 independent observations from empty chambers across 3 experimental years, which was used to determine the sensitivity of the platform to measure pulmonary gas exchange from animals with very low lung volume (pups less than 4 days of age) or septic pups. For experiments involving collecting Promethion readings at 30°C, RC3 chambers were placed in a Datesand Maxi Thermacage (4 chambers per cage), with the temperature set to 30°C. The Thermacages were turned on for at least 10 minutes prior to placing pups within RC3 chambers.

**Figure 1.**
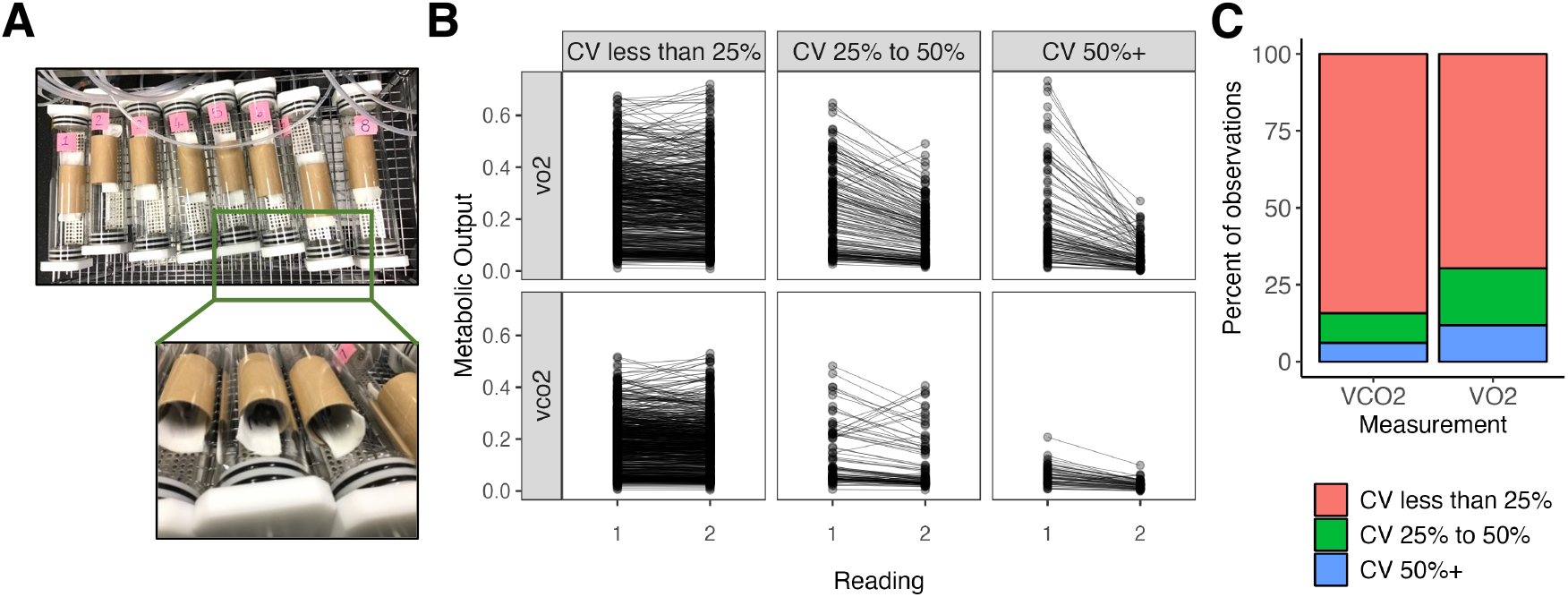
RC3 chamber configuration and value variance between reading 1 and reading 2 across ten different experiments performed between 2021 and 2023. These data represent 29 measurements from empty tubes and 1584 animal measurements across nine different experiments. A. RC3 chamber setup, depicting numbered tubes with cardboard rolls and wipes. B. Promethion-RC3 chamber coefficient of variance by replicate readings from VO2 and VCO2 values. C. Percentage of observations within each CV threshold for VO2 and VCO2

### Data processing

The raw data collected during the experiment consisted of two main components: a wide-format metadata file containing information about the experimental animals (e.g., animal IDs, cage assignments, dates of birth, sex) and metabolic data from the Promethion metabolic monitoring system. Raw data files include the rate of oxygen consumption in ml/min (VO2) and rate of carbon dioxide emission in ml/min (VCO_2_).

Promethion data analysis software package, MacroInterpreter (version 2.48) by Sable System, was used to conduct primary data processing of fundamental metabolic transformations. Sable System provided a custom-made Macro (OneClickMacroV.2.51.2-slice3mins) to handle fundamental metabolic calculations and generate the necessary metabolic data variables for our analyses, such as respiratory quotient (RQ) and kilocalories per hour (Kcal/hr), based on the raw VO2 and VCO2 values generated by the Promethion Core CGF system for each cage. The final output of executing the Macro is a .csv file with data in a format that can easily be used programmatically. The Promethion Core CGF software generates one data file per cage per time point. For an experiment consisting of 10 time points and 4 cages, 40 separate csv files with raw values are generated that need to be bulk-processed via MacroInterpreter and exported in a final usable format. To facilitate data processing, we created custom scripts to combine these individual experimental files and merge them with other experimental variables (e.g. animal sex, treatment group). This template is available in the supplement. Prior to biological analysis, all animal measurements with a coefficient of variation (CV) over 50% for either VO2 or VCO2 were removed. For empty cages, outlier values (values beyond the 97.5^th^ percentile of all measurements taken) were also removed, as these measurements were likely mis-labelled as empty while they indeed contained pups.

### Statistical analysis

All data exploration, visuals and statistics were conducted using R version 4.3.2. Statistical analyses were performed using base R functions, and graphics produces using ggplot2. The log-rank test was used to compare survival probability between experimental groups. The Wilcoxon rank-sum test was used to perform pairwise comparisons of VO2, VCO2, and temperature variables. The Fisher’s exact test was used to compare the proportions for readings with high CV values.

### Data availability

All data and analytical scripts presented in this manuscript are publicly available on GitHub: https://github.com/ImmuneResilience/ic_preclin_neonatal_methods.

## Results

### Cross-sectional IC measurements display low coefficients of variance

Given that our analytical strategy involves averaging out two measurements per experimental animal for each time point, we first investigated the coefficient of variance (CV) observed across all experimental measurements collected from 2021 to 2023. These data demonstrated that for VO2, 84% of observations held a CV below 25%. For VCO2, 69% of observations fell below 50% (Fig. 1B-C). A smaller proportion of observations were found to vary by more than 50%, and these were also driven by pups whose first values were dramatically higher than their 2^nd^ value (Fig. 1B). For example, for readings with a CV less than 25%, there are equal proportions of instances where reading 1 is higher than reading 2, or vice versa. However, readings where the CV was higher than 25% were almost exclusively those where the first reading was higher than the 2^nd^ (Fig. S1). Of note, readings with high CVs were not enriched according to the pups’ age or sickness. Upon analysis of the experimental metadata we identified that readings with a CV over 50% occurred exclusively during the first experimental year (2021), when regular machine maintenance had been postponed due to COVID-19. Following the re-establishment of yearly maintenance, significantly fewer instances of variable readings were observed (Fig. S1C), suggesting that the initial measurements were in part due to machine error. Based on these data, we removed all observations for VO2 and VCO2 with a CV higher than 50%.

### Sensitivity of RC3 chambers for very young or sick neonatal pups

To investigate the biological limit of RC3 chambers to measure VO2 and VCO2 values from animals with extremely low pulmonary capacity (neonatal mice younger than DOL 8) or with a low respiratory rate (septic neonates), we compared VO2 and VCO2 measurements from pups on the first to 7^th^ day of life and 7-day-old pups challenged with polymicrobial sepsis compared to healthy 7-day-old pups. VO2 was reliably captured from DOL1 onwards, where only one observation fell within the range of empty tubes on DOL1, and all measurements were within the detectable range thereafter. For VCO2, less than 10% of measurements (two data points) fell within the range of empty tubes. (Fig. 2A-B). To determine the sensitivity to capture VO2 and VCO2 among septic mice, we compared healthy 7-day-old pups with septic pups 1-3 hours post challenge, the time when we observe the lowest VO2 and VCO2 values among sick, challenged pups. All but one sick pup had readings above the baseline, suggesting that this platform is sensitive enough to capture gas exchange even when lung volume is limited or respiratory rate depressed.

**Figure 2.**
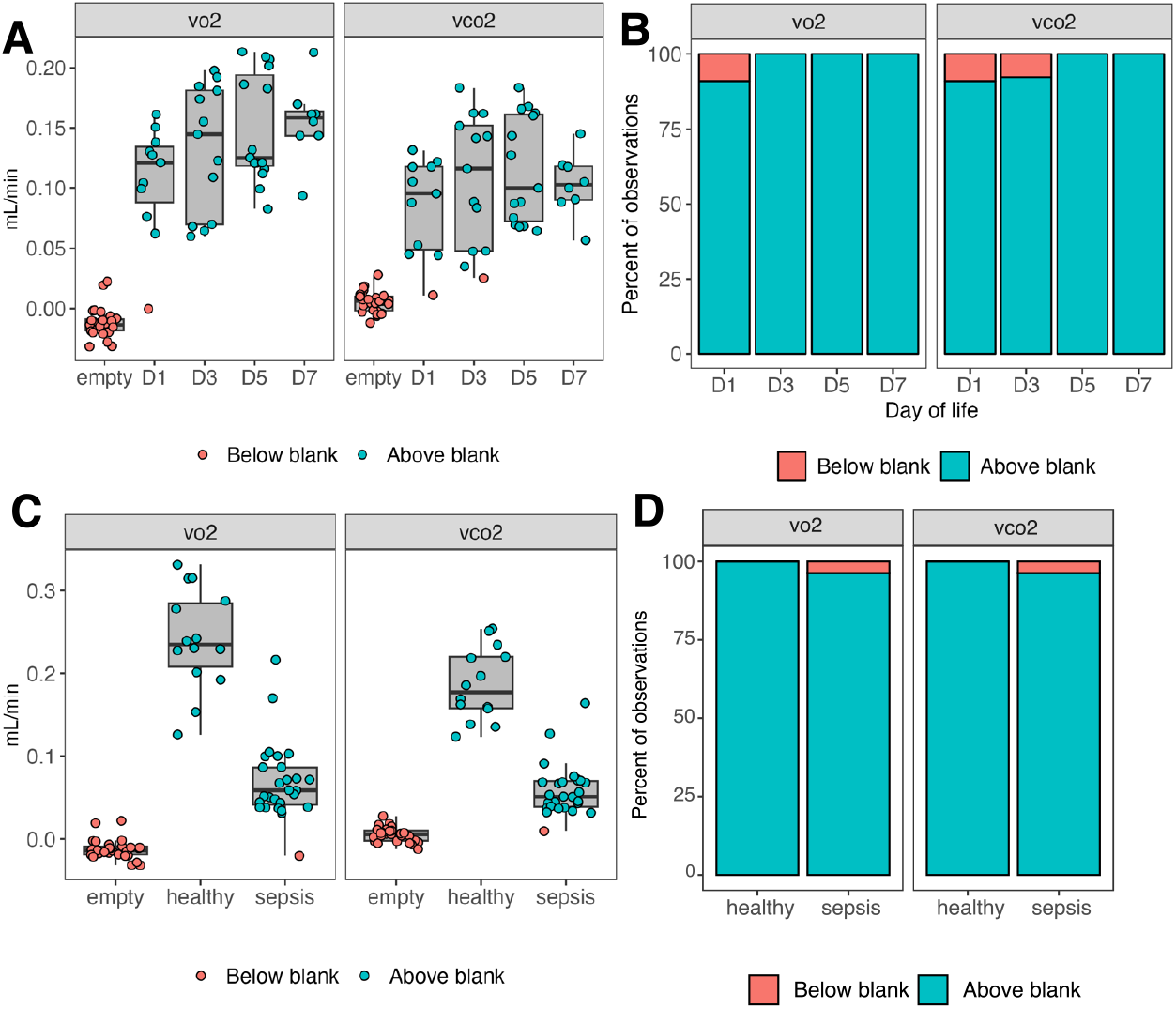
Promethion-RC3 chamber sensitivity among very young or sick neonatal mice. A-B. VO2 and VCO2 among 1-7 day-old pups and empty RC3 chambers. A. Absolute VO2 and VCO2 values. B. Percent of values at each time point that fall below the maximum values from empty chambers. n = 8-15 pups per time point, 21 observations for empty chambers. C-D. VO2 and VCO2 of septic or healthy 7-day-old pups compared to empty chambers. C. Absolute VO2 and VCO2 values. D. Percent of observations among health or septic pups that fall below the maximum values of empty tubes. n = 14 observations for healthy, 36 observations for septic mice, and 21 observations for empty chambers

### Use of the Promethion Core does not significantly alter outcomes during sepsis

Given the additional animal handling involved in measuring the metabolic status of pups compared to the normal monitoring schedule used for challenge experiments, it was possible that the stress that animals were exposed to during this procedure could alter the sepsis response and render animals more susceptible to reaching humane endpoint. These stressors involve more frequent separation from the dam at early time points post challenge and increased separation time from the dam (average of 14 minutes for visits involving IC measurements compared to 5 minutes for standard welfare monitoring) for each visit. Removal from the nest also deprives the pups of warmth received from the dam or littermates. To determine if this could be the case, we challenged four experimental litters with an LD30 of cecal slurry, and then randomized each cage to either the Promethion experimental schedule or no Promethion, where the standard monitoring practice was used (Figure 3A). While there was a trend for some pups to reach humane endpoint earlier among the Promethion compared to the No Promethion pups, the difference in overall survival between groups was not significant (Fig. 3B). Among pups that did not reach humane endpoint, we observed an overall increase in change in weight from baseline for pups assigned to the Promethion group after approximately 20 hours post challenge (Fig. 3C). Distribution of clinical scores was similar between groups at baseline, 20, and 48 hours post challenge. More variance in clinical scores was observed among the Promethion group at 12 hours post challenge than for the No Promethion group (Fig. 3D). Overall, this suggests that the impost associated with intensive early measurements does not render pups more susceptible to polymicrobial sepsis.

**Figure 3.**
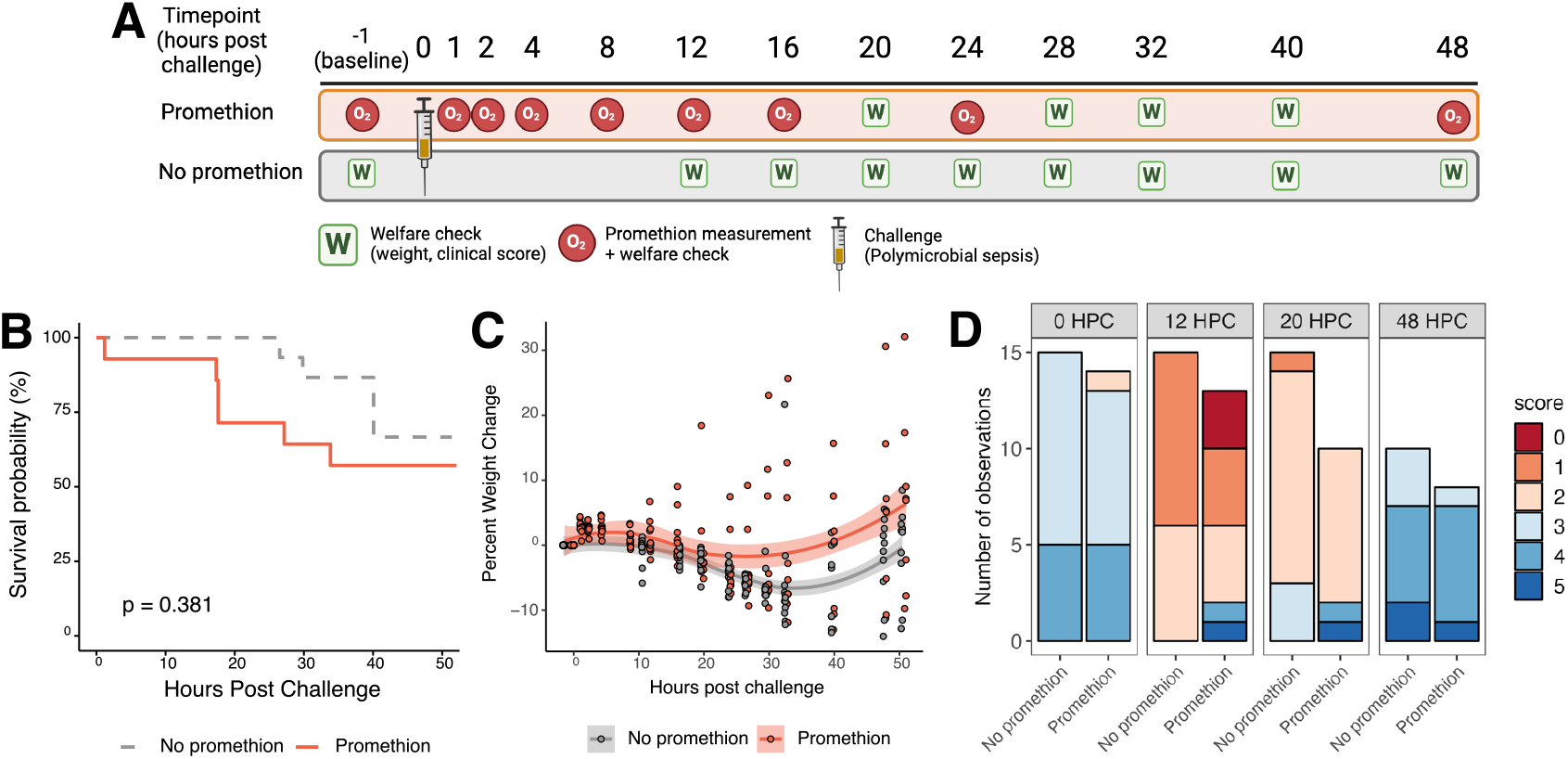
Impact of IC-associated impost on polymicrobial sepsis outcomes. A. Experimental design, indicating timing of challenge, welfare checks, and promethion measurements. B. Overall survival of pups assigned to Promethion or no Promethion monitoring schedules (p = 0.381, log-rank test). C. Percent weight change from time of challenge among pups assigned to Promethion or No promethion monitoring schedules. Shaded area indicates 95% confidence interval. D. Clinical scores at 0, 12, 20, and 48 hours post challenge for pups assigned Promethion or No promethion monitoring schedules. n = 15 Promethin, 14 No promethion. Panel A was created with Biorender.

### Performing Promethion measurements at 30°C compared to room temperature minimizes animal heat loss and has a minor impact on VO2 and VCO2 readings

While the impost associated with intensive Promethion readings did not significantly worsen sepsis outcomes, we did observe that external body temperature decreased by about 5°C during the time spent in the RC3 chamber. Given the importance of thermoregulation on animal welfare, we still sought to determine whether performing the readings at 30°C (the approximate temperature within the nest) would lessen heat loss, and if VO2 and VCO2 readings differed between these conditions. To compare IC readings at 30°C and room temperature (RT), we placed four of eight RC3 chambers in a ThermoCage set to 30°C, and left the other four chambers at RT. We then randomized pups within four litters to either the 30°C or RT chambers and challenged two of the four cages with cecal slurry LD10 dose (Fig. 4A). Pup temperatures were measured after the dam was removed from the cage (temp) and after the pups were removed from the RC3 chambers (temp2). While the mean difference in temperature was 0.98°C for the pups in the 30°C condition across all time points, pups in the RT condition lost on average 4.4°C, consistent with our previous observations (Fig 4B-C).

**Figure 4.**
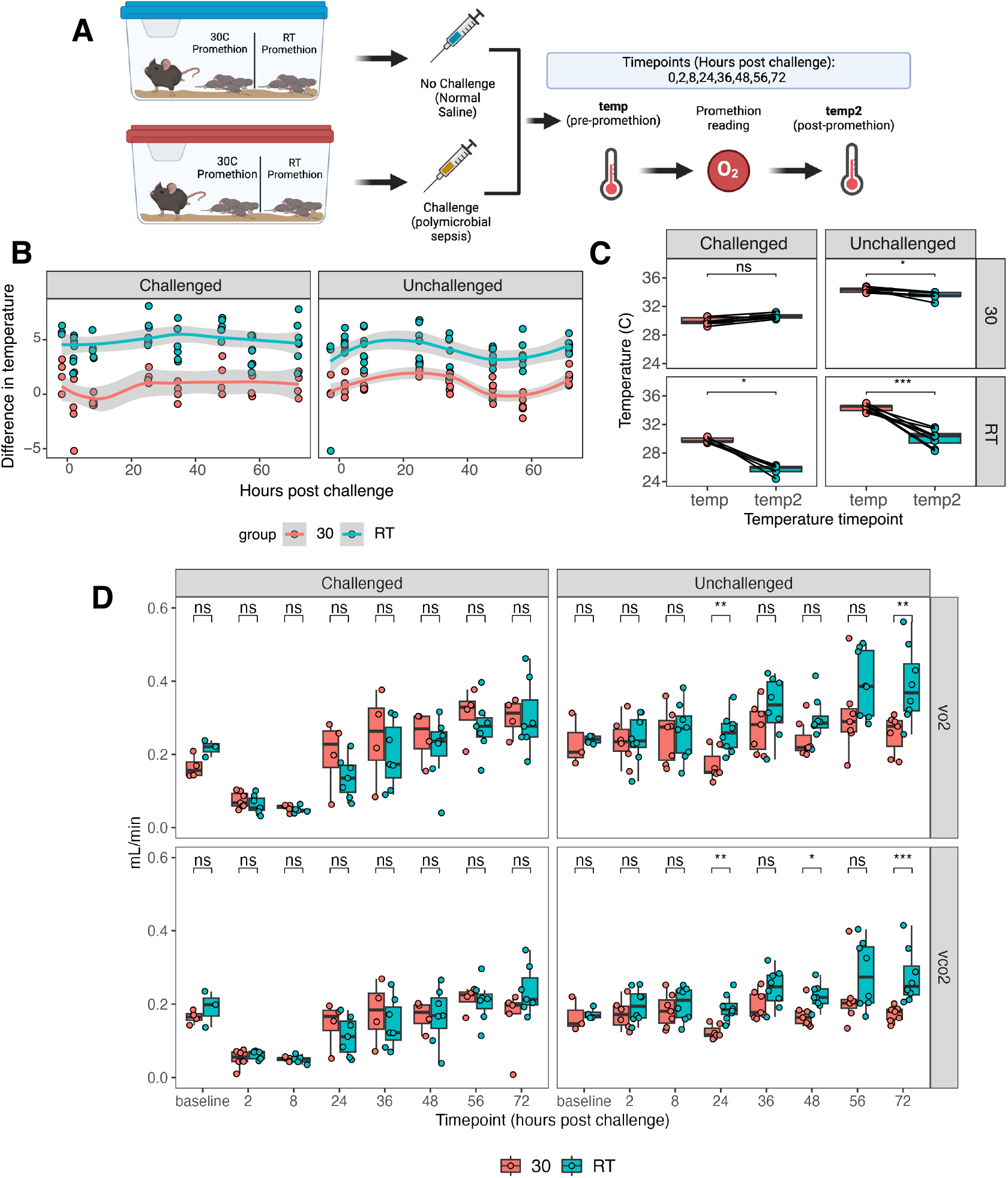
Impact of RC3 chamber temperature on animal temperature loss and gas exchange. A. Experimental design, indicating within-cage randomization to 30C or room temperature (RT) Promethion readings, combined with between-cage randomization of sepsis vs. control pups. B. Difference in pup temperature from the 2^nd^ to the first temperature reading. Shading indicates 95% confidence C. Paired observations at eight hours post challenge, the lowest body temperature observed among septic pups, showing minimal changes in temperature among 30°C pups and consistent drops in temperature for both challenged and unchallenged pups in the RT condition. D. Comparison of VO2 and VCO2 readings taken and baseline, and then 2-72 hours post challenge. ^*^ p < 0.05, ^**^ p < 0.01, ^***^ P < 0.001, Wilcoxon Rank-Sum test. 30°C n = 8 challenged, 7 unchallenged; RT n = 8 challenged, 7 unchallenged. Panel A was created using Biorender.

We observed differences in VO2 and VCO2 between the conditions, for unchallenged mice only, whereby at room temperature pups had higher and more variable VO2 and VCO2 values after DOL8 (24 hours post challenge). While the coefficients of variation for these measurements were mostly less than 25%, we observed that VO2 and VCO2 values increased from the first to the second replicate for most of the animals at the RT condition, while the opposite trend was seen for the 30C condition by the time the animals reached DOL10 (72 hours post challenge). Furthermore, this relationship was not observed at earlier time points (Suppl. Fig. 2). This suggests that the temperature of RC3 chambers in older mice may differentially impact the metabolic rate of healthy animals, and the implications of this difference should be considered in the experimental design.

## Discussion

Here, we applied a custom configuration of the Promethion Core System with RC3 chambers to determine the VO2 and VCO2 of healthy and septic neonatal pups from day of life 1 to 7. We found this experimental platform adapted to newborn mice, i) delivers stable values (low CV) across repeat measurements; ii) efficiently captures gas exchange even in very young (day of life 1) and/or very sick (septic) mice; and iii) does not impact health outcomes in a neonatal sepsis model, especially if body temperature of the newborn animal is supported during handling. This platform thus appears suitable for longitudinal indirect calorimetry assessments in newborn mice.

Given our focus on neonatal sepsis, we sought to determine whether our custom configuration of the Promethion Core System with RC3 chambers was sensitive enough to measure the VO2 and VCO2 of either very young, or sick animals whose pulmonary gas exchange rates are low due to either limited lung volume for young mice, or a hypometabolic state and low respiratory rate for septic animals. Compared to a range of blank values collected across multiple experiments from 2021-2023, we were able to measure VO2 values for almost all experimental animals. However, especially for sick pups, the values we observed were close to the range blank values. While the proportion of these measurements were exceedingly small, experimental error is possible in this range and individual teams will need to decide how to handle these data points.

Use of the Promethion chamber early during the septic disease course would provide valuable insight into the early physiological adaptations to infection as in our experimental model more than half of septic neonates reach humane endpoint within the first three days. This is at a time when the animals appear physiologically similar, and where sample harvests would necessitate sacrificing the animals. While our approach to IC is non-invasive, it was still possible that the frequent handling, the time away from the dam, isolation from littermates, with additional cold exposure could alter the disease course. Yet, we did not identify a significant impact including IC on sepsis severity measured by overall survival, degree of weight loss, and clinical score. It may have been plausible for the additional cold exposure to have increased survival in some settings, as studies have shown that reducing body temperature in adult rodents improves survival probability in experimental models of ischemia and hypoxia, for example (*18*). Thus, our findings with the neonatal polymicrobial sepsis model may not apply to other experimental models. It is also possible that the use of the Promethion chamber may render pups more susceptible to other experimental challenges. For example, while frequent handling increased the ability of murine pups to respond better to acute stress and to adapt to the environment, it also resulted in dysfunction of the reproductive, renal, social and neural pathways, which leads to chronic behavioural deficits (*19*).

Minimizing temperature loss during the process of collecting indirect calorimetry data may further improve animal welfare, and minimize possible artefacts introduced by the Promethion system. Body temperature for both healthy and septic pups decreased by an average of 4°C from the time the dam was removed from the cage, to when the pups were removed from the Promethion Chamber. This finding is consistent with thermal adaptation observed among common house mice pups that, when isolated from the nest and placed in ambient temperature, decrease their core temperature to just slightly above the ambient temperature (*20*). Generally, in laboratory adult mice, the housing temperature (20-24ºC), when lower than the mouse’s critical temperature, increases metabolism to counter the heat loss at approximately 30°C (*21, 22*). Mice being an altricial species and mice pups at birth being ectothermic cannot thermoregulate (regulate their body temperature) for the first 2-3 weeks of life (*23, 24*) and rely on the heat from the dam, littermates in the nest and heat retained by the nest itself for survival (*25-28*). To limit the effect of the Promethion on loss of body heat, we placed the RC3 chambers in ThermoCages heated to 30°C. This refinement was successful in limiting change in body temperature from the time pups were separated from the dam to when they were removed from the RC3 chambers. Importantly, pups normally became hypothermic for 12-24 hours after challenge, measurable within 2 hours of challenge (Fig. 4B), which is also consistent with the hypothermic response to sepsis observed among adult rodents (*13*). In the 30°C condition, septic pups that were hypothermic were not excessively warmed by this approach, as their body temperature was maintained, but not increased, by being placed in the 30°C chambers, and was still approximately 4°C cooler compared to healthy pups. This is reassuring, as it suggests the additional heat did not alter the hypothermic response, but appeared to only replace the heat that would have otherwise been provided by the dam within the nest. The caveat to this approach is that, after DOL8, when the animals are furred and are able to thermoregulate, the heated chambers may alter the metabolic adaptation of animals placed within the chambers. The Promethion readings performed between DOL 1-7 and at 30°C (Fig. 2), demonstrated an increase in VO2 and VCO2 as the pups grew older. The lack of increase in VO2 and VCO2 for animals housed at 30°C between DOL 7 and 9 in the 30°C condition was surprising, especially since at room temperature, this age-dependent increase was still observed. It thus is possible that warmed chambers are suitable for neonatal animals in the first week of life, but room temperature readings preferrable for older animals. With this, animal health status and age should be considered when selecting RC3 chamber temperatures. Very young pups without fur or sick pups may require additional warmth, while older and furred pups may require RC3 chambers to be kept at room temperature.

We present here the raw data acquired from the Promethion chamber. However, analytical decisions still need to be made for teams employing this strategy. For example, for VO2 and VCO2 levels that fall within the range of empty chambers, these values can either be censored, floored to a constant value, or left unaltered. These decisions would impact some analyses more than others. For example, when for comparing of hypometabolic to normometabolic animals, where the difference between groups is dramatic, leaving the low values unaltered is unlikely to alter the conclusions of the experiment. However, if comparing one group of hypometabolic animals to another, then some values derived from the VO2 and VCO2 (e.g. the respiratory quotient) may potentially be impacted depending on how the low values are handled. Accurate reporting and regular data collection of empty tubes ensures that any limitations can be rigorously evaluated and assessed. Lastly, we identified that, by collecting two measurements per animal per time point, up to 13% of measurements displayed high variance that was driven mostly by an initial value dramatically higher than the 2^nd^ value. We identified these variable measurements in early experimental years but not later once annual maintenance was reinstated. Thus, it is possible that even fewer measurements would need to be removed when regular servicing is maintained. Irrespective of the data handling, to ensure the reliability of the system, monthly calibrations of the gas sensors are thus recommended as well as yearly maintenance by a qualified technician.

The evaluation of RC3 chambers for neonatal pups could be improved in the future by, for example, determining the stability of cross-sectional readings beyond the two cycles we used for our studies. We also did not attempt measurements on DOL 0. However, given the range of values that can be captured on DOL 1, it is possible that even on DOL 0, VO2 and VCO2 can be captured for most pups. While extended periods of isolation from the dam and littermates can potentially create artefacts, it would also be worthwhile to determine if measurements beyond the two readings provide a more accurate estimate. Further, the utility of taking whole-litter measurements by removing the dam from the cage can potentially provide useful insight at the whole-litter level. While such an approach would fail to provide measurements for individual pups, it may be appropriate in circumstances where treatment allocations for animals are done across cages, and not within litters. Lastly, given that the Promethion Core is most commonly used for longitudinal measurements, the current software is not well designed to handle cross-sectional data, and the export of individual files is time consuming. While our team developed extensive custom scripts to then merge these various files, there is an opportunity to develop software to streamline this process for researchers without requiring teams to write custom pipelines.

Overall, the Promethion-RC3 configuration represents a valuable tool to determine whole body metabolism at an individual level for very young or sick murine pups. The low level of impost, especially when using warmed RC3 chambers for young animals, provides an excellent opportunity to integrate this non-invasive tool to longitudinally determine host metabolic status in the neonatal period.

## Supplementary Figures

**Supplementary Figure 1.**
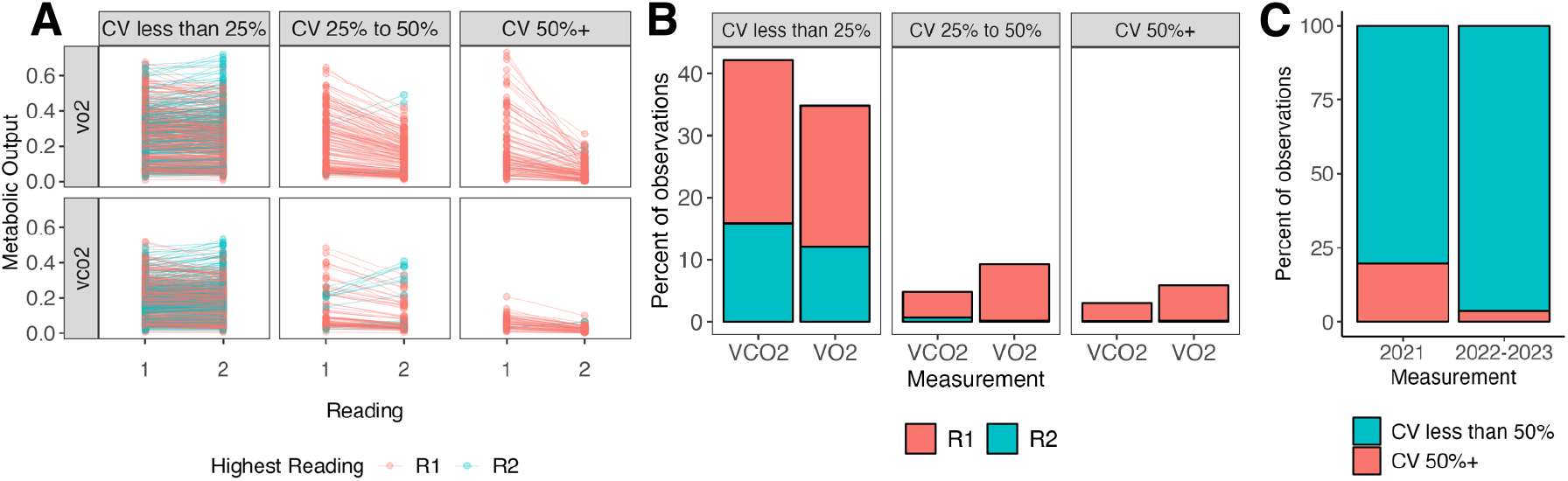
Difference between reading 1 and reading 2 in relation to CV. A. Paired readings stratified by the CV. B. Percentage of readings within VO2 and VCO2 where reading 1 (R1) or reading 2 (R2) was the highest. C. Proportion of readings with a CV over 50% across experimental years significantly differed between 2021 vs. 2022 – 2023 (p < 0.05, Fisher’s Exact Test).

**Supplementary Figure 2.**
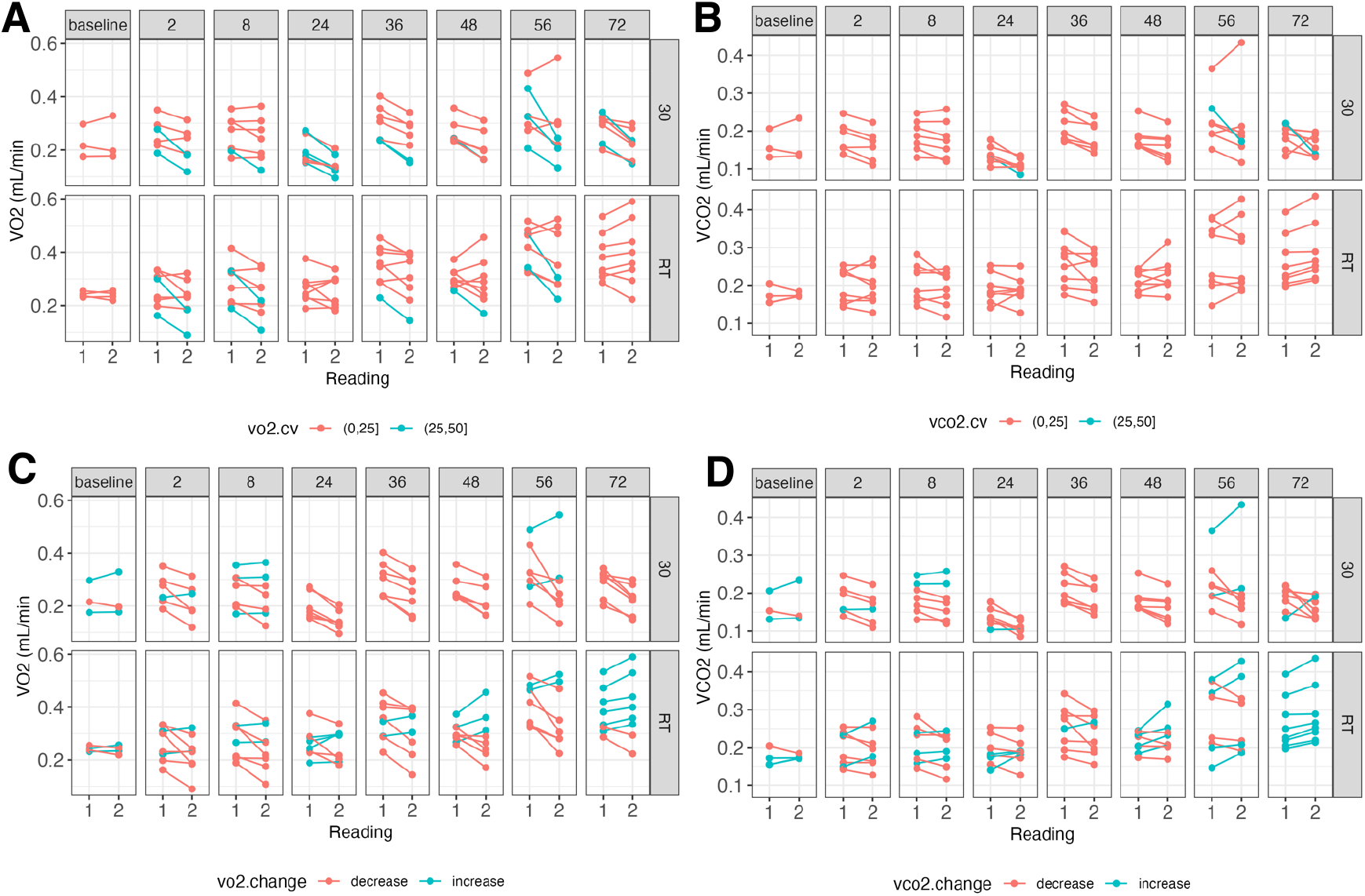
Technical Promethion replicates for pups randomized to readings at 30°C or room temperature. A-B. VO2 technical replicates, colors indicating (A) coefficient of variation either between 0 and 25% or 25% to 50% or B. whether the 2^nd^ reading increased or decreased relative to the first reading. C-D. VCO2 technical replicates, colors indicating (A) coefficient of variation either between 0 and 25% or 25% to 50% or B. whether the 2^nd^ reading increased or decreased relative to the first reading.

## Notes

https://github.com/ImmuneResilience/ic_preclin_neonatal_methods

### Competing Interest Statement

The authors have declared no competing interest.

## References

1. A. Wang, H. H. Luan, R. Medzhitov, An evolutionary perspective on immunometabolism. Science 363, (2019).

2. J. Ye, R. Medzhitov, Control strategies in systemic metabolism. Nat Metab 1, 947–957 (2019).

3. A. Wang, R. Medzhitov, Counting Calories: The Cost of Inflammation. Cell 177, 223–224 (2019).

4. J. L. Wynn et al., Increased mortality and altered immunity in neonatal sepsis produced by generalized peritonitis. Shock (Augusta, Ga.) 28, 675–683 (2007).

5. M. Delsoglio, N. Achamrah, M. M. Berger, C. Pichard, Indirect Calorimetry in Clinical Practice. J Clin Med 8, (2019).

6. H. Henneicke et al., Skeletal glucocorticoid signalling determines leptin resistance and obesity in aging mice. Mol Metab 42, 101098 (2020).

7. S. M. Solon-Biet et al., Branched chain amino acids impact health and lifespan indirectly via amino acid balance and appetite control. Nat Metab 1, 532–545 (2019).

8. M. A. McCann et al., Adipose expression of CREB3L3 modulates body weight during obesity. Scientific reports 11, 19400 (2021).

9. D. Binyamin et al., The aging mouse microbiome has obesogenic characteristics. Genome Med 12, 87 (2020).

10. B. Zhou et al., Serum- and glucocorticoid-induced kinase drives hepatic insulin resistance by directly inhibiting AMP-activated protein kinase. Cell reports 37, 109785 (2021).

11. D. Wang et al., LSD1 mediates microbial metabolite butyrate-induced thermogenesis in brown and white adipose tissue. Metabolism 102, 154011 (2020).

12. C. C. Chiang, M. Korinek, W. J. Cheng, T. L. Hwang, Targeting Neutrophils to Treat Acute Respiratory Distress Syndrome in Coronavirus Disease. Front Pharmacol 11, 572009 (2020).

13. K. Ganeshan et al., Energetic Trade-Offs and Hypometabolic States Promote Disease Tolerance. Cell 177, 399–413.e312 (2019).

14. K. J. Cummings, C. Willie, R. J. Wilson, Pituitary adenylate cyclase-activating polypeptide maintains neonatal breathing but not metabolism during mild reductions in ambient temperature. Am J Physiol Regul Integr Comp Physiol 294, R956–965 (2008).

15. B. Brook et al., A Controlled Mouse Model for Neonatal Polymicrobial Sepsis. Journal of visualized experiments: JoVE, (2019).

16. B. Brook et al., Robust health-score based survival prediction for a neonatal mouse model of polymicrobial sepsis. PloS one 14, e0218714 (2019).

17. B. Brook et al., BCG vaccination-induced emergency granulopoiesis provides rapid protection from neonatal sepsis. Science translational medicine 12, (2020).

18. J. G. Christopher, The therapeutic potential of regulated hypothermia. Emergency Medicine Journal 18, 81 (2001).

19. C. Raineki, A. B. Lucion, J. Weinberg, Neonatal handling: an overview of the positive and negative effects. Developmental psychobiology 56, 1613–1625 (2014).

20. M. S. Blumberg, in Developmental Psychobiology, E. M. Blass, Ed. (Springer US, Boston, MA, 2001), pp. 199–228.

21. F. Andrews, Temperature regulation in laboratory rodents: Christopher J. Gordon. Cambridge University Press, 1993. ISBN 0-521-41426-1 hardback. Journal of Thermal Biology 20, 365–366 (1995).

22. B. N. Gaskill et al., Heat or Insulation: Behavioral Titration of Mouse Preference for Warmth or Access to a Nest. PloS one 7, e32799 (2012).

23. R. J. Berry, F. H. Bronson, Life history and bioeconomy of the house mouse. Biological Reviews 67, 519–550 (1992).

24. R. H. Ogilvie Dm Fau-Stinson, R. H. Stinson, The effect of age on temperature selection by laboratory mice (Mus musculus).

25. B. N. Gaskill et al., Impact of nesting material on mouse body temperature and physiology. Physiology & Behavior 110-111, 87–95 (2013).

26. B. N. Gaskill et al., Energy Reallocation to Breeding Performance through Improved Nest Building in Laboratory Mice. PloS one 8, e74153 (2013).

27. N. Latham, G. Mason, From house mouse to mouse house: the behavioural biology of free-living Mus musculus and its implications in the laboratory. Applied Animal Behaviour Science 86, 261–289 (2004).

28. E. M. Weber, I. A. S. Olsson, Maternal behaviour in Mus musculus sp.: An ethological review. Applied Animal Behaviour Science 114, 1–22 (2008).

